# q-mer analysis: a generalized method for analyzing RNA-Seq data

**DOI:** 10.1101/2021.05.01.424421

**Authors:** Tatsuma Shoji, Yoshiharu Sato

**Author notes:** (/+81-03-5777-1689).

## Abstract

RNA-Seq data are usually summarized by counting the number of transcript reads aligned to each gene. However, count-based methods do not take alignment information, where and how each read was mapped in the gene, into account. This information is essential to characterize samples accurately. In this study, we developed a method to summarize RNA-Seq data without losing alignment information. To include alignment information, we introduce “q-mer analysis,” which summarizes RNA-Seq data with 4q kinds of q-length oligomers. Using publicly available RNA-Seq datasets, we demonstrate that at least q ≧ 9 is required for capturing alignment information in *Homo sapiens*. Furthermore, principal component analysis showed that q-mer analysis with q = 14 linearly distinguished samples from controls, while a count-based method failed. These results indicate that alignment information is essential to characterize transcriptomics samples. In conclusion, we introduce q-mer analysis to include alignment information in RNA-Seq analysis and demonstrate the superiority of q-mer analysis over count-based methods in that q-mer analysis can distinguish case samples from controls. Combining RNA-Seq research with q-mer analysis could be useful for identifying distinguishing transcriptomic features that could provide hypotheses for disease mechanisms.

## 1. Introduction

RNA-Seq is commonly used in molecular biology [1] since its development more than a decade ago [2–6]. Many studies have characterized samples at the transcriptional level using RNA-Seq or at the post-transcriptional level by coupling the appropriate biochemical assay with RNA-Seq [7-9]. The number of publications containing RNA-Seq data is approximately 36,000 in 2021 (PubMed). Furthermore, many types of software have been developed for analyzing RNA-Seq data [10–12]. RNA-Seq is now an indispensable technology.

In terms of RNA-Seq data analysis, the core task of mapping and counting reads is common to many kinds of software. After the mapping process, RNA-Seq data is summarized by counting the reads aligned to each exon, gene, or transcript [11] and is reported in the form of gene expression tables [13–16]. However, count-based methods do not consider alignment information. In other words, count-based methods discard information regarding where and how the aligner mapped each read in the exon, gene, or transcript. If the difference between samples and controls is well characterized at the transcriptional level, and differential gene expression analysis is sufficient for the data analysis, count-based methods work well [17]. There is a valid statistical theory that explains the fluctuations of the read counts [18–21], and many software products test whether the observed difference between samples and controls is significant [19–23]. However, if the aim of the study involves post-transcriptional regulation or mutations in the genome, the loss of alignment information is problematic for two reasons. First, count-based methods ignore mismatches in alignment information, which are “expressed mutations,” and thus are critical features of samples. Second, count-based methods do not take the number of reads aligned to the transcriptome at a unique region into account, which retains post-transcriptional information: the secondary or tertiary structure of mRNA or protein–mRNA interactions, including the ribosomal distribution across mRNAs [24], which potentially reflects the sample characteristics of interest [25]. Therefore, RNA-Seq data should be summarized in a more informative way by taking alignment information into account to accurately describe samples.

Although several studies have described the usefulness of alignment information and used it to detect mRNA isoforms [17,26], applications for taking alignment information into account are limited. Currently, no studies focus on summarizing alignment information in RNA-Seq data.

In this study, we provide a method to describe samples more accurately by including alignment information. First, we introduce a new method called “q-mer analysis” that summarizes RNA-Seq data without losing alignment information using a “q-mer vector,” which is the occurrence rate of 4^q^ kinds of q-length oligomers in RNA-Seq data. We then determine the appropriate q value and assess the ability of q-mer analysis to describe samples using two publicly available RNA-Seq datasets from Homo sapiens. Further investigation shows the superiority of q-mer analysis to count-based methods.

## 2. Results

### 2.1. q-mer Analysis of RNA-Seq Data

The simplest way to approximate RNA-Seq data is using the ratio of A, T, G, and C nucleotides in the alignment data. The second simplest way to approximate the RNA-Seq data is using the ratio of the 4^2^ kinds of oligomers, namely, AA, AT, AG, AC, TA, TT, TG, TC, GA, GT, GG, GC, CA, CT, CG, and CC, in the alignment data. In this way, we can approximate the RNA-Seq data and express alignment information at the same time using the occurrence rate of 4^q^ kinds of q-length oligomers (Figure 1). If q is large enough, this approximation includes count-based RNA-Seq data. Hereafter, we introduce this approximation as “q-mer analysis” and the resulting 4^q^ dimension vector and matrix as the “q-mer vector” and “q-mer matrix,” respectively.

**Figure 1.**
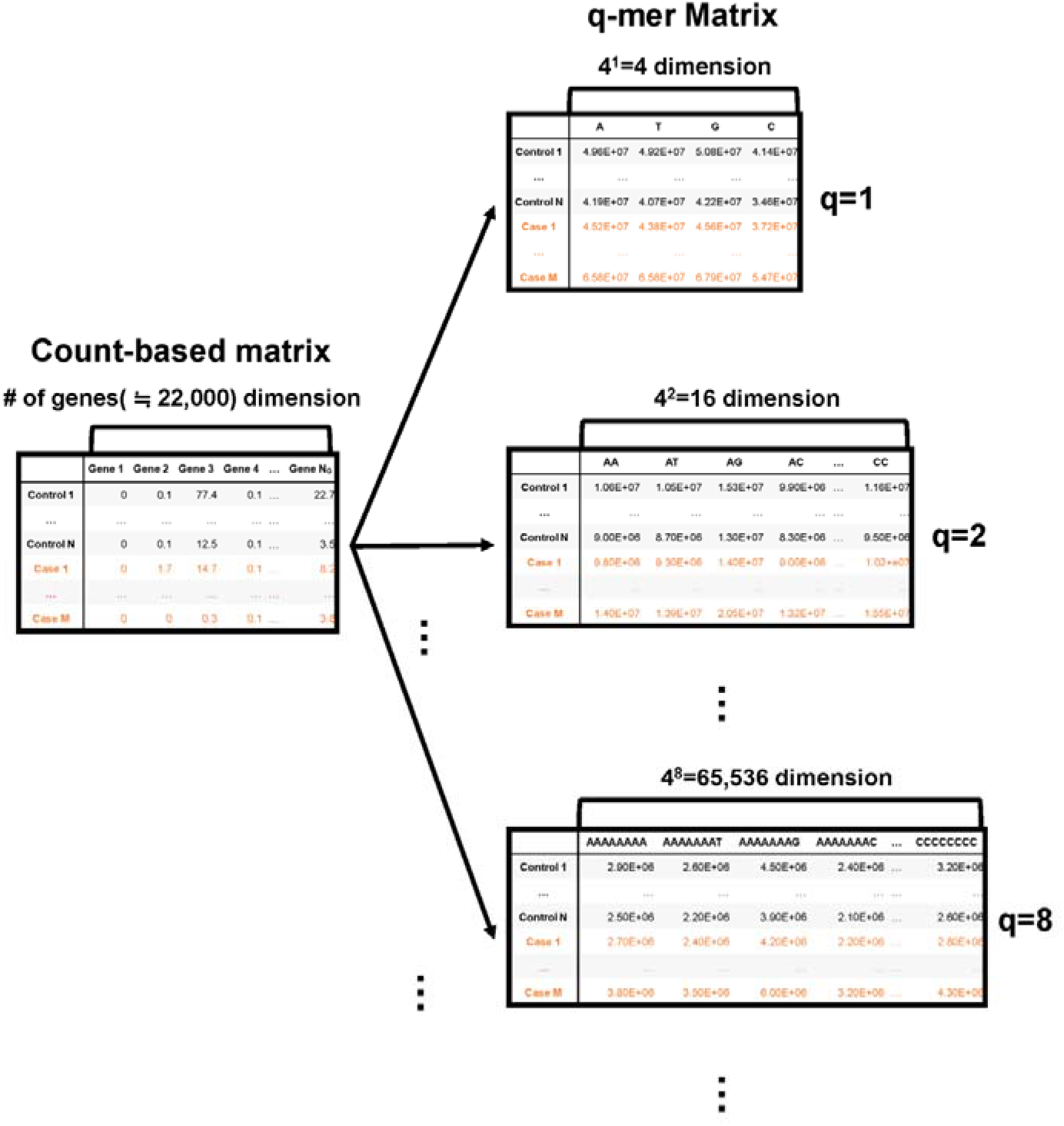
Schematic image of the count-based matrix and the q-mer matrices. The count-based summary of RNA-Seq data is shown on the left, while q-mer matrices are shown on the right. The number of the explanatory variables is shown above each table. The q value is shown to the right of each table.

### 2.2. q ≧ 9 is Required to Express Alignment Information in H. sapiens

To investigate the appropriate q value to express alignment information of RNA-Seq data in *H. sapiens*, we calculated the q-mer vector with a q value of 1 to 14 for two publicly available RNA-Seq datasets from *H. sapiens* (GEO accession numbers GSE99349 [27] and GSE106589 [28]). The first RNA-Seq experiment (GSE99349) was performed on neuronal nuclei isolated from the post-mortem dorsolateral prefrontal cortex of 19 cocaine-addicted and 17 healthy control cases. The second RNA-Seq experiment (GSE106589) was performed on 18 human-induced pluripotent stem cell-derived neural progenitor cells (hiPSCs-NPCs) derived from 14 individuals with childhood-onset schizophrenia (COS) and 20 hiPSCs-NPCs from 12 unrelated healthy controls. Regarding the selection of the RNA-Seq datasets, see the selection criteria described in Section 4. The rationale for picking a range of q values from 1 to 14 is that the total length of all mRNAs in *H. sapiens* (589,150,963) is larger than 4^14^ but less than 4^15^, which means the q-mer vectors with q values greater than 14 are sparse.

Table 1 summarizes the number of zero elements observed in the q-mer vectors in the example data. Surprisingly, zero elements were not observed in q-mer vectors with q values less than 10, indicating that q = 9 is at least required to express alignment information from RNA-Seq data in *H. sapiens*. Note that 4^9^ (262,144) is approximately 10 times the number of genes in *H. sapiens*, suggesting that the dimensions of gene expression tables are too low to accurately summarize RNA-Seq data.

**Table 1.**
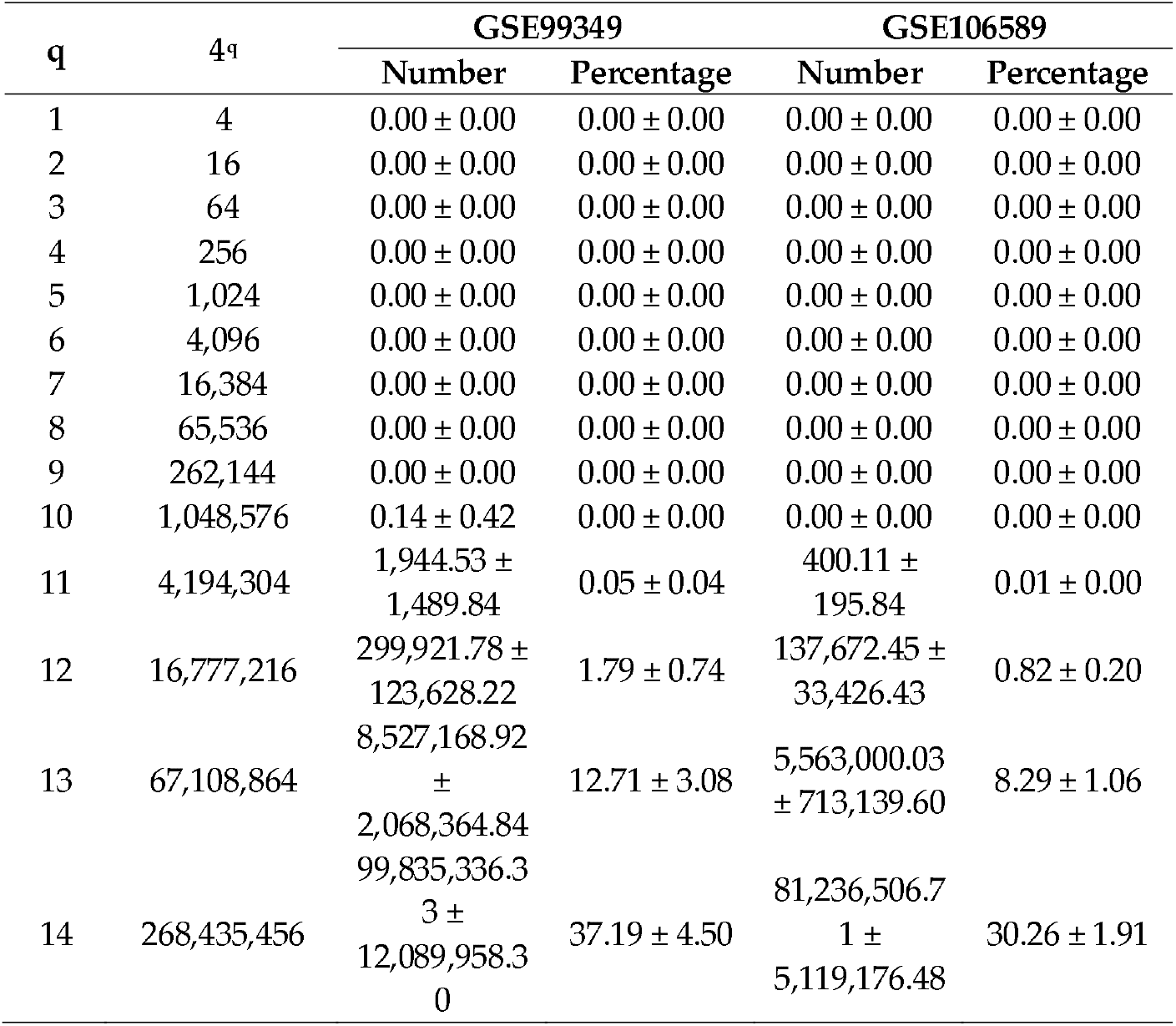
The sparsity of q-mer vectors in *H. sapiens*. The mean and S.D. of the number or percentage of zero elements in the q-mer vectors are shown in the “Number” or “Percentage” columns, respectively.

### 2.3. Sample Description with q-mer Analysis

The high dimensionality of RNA-Seq data in *H. sapiens* may result from the inclusion of alignment information. This very large number led us to speculate that q-mer analysis could describe the samples more accurately than could count-based methods. To investigate the ability of q-mer analysis to describe these samples, we applied principal component analysis (PCA) to the two RNA-Seq datasets described above after preprocessing (Figure 2).

**Figure 2.**
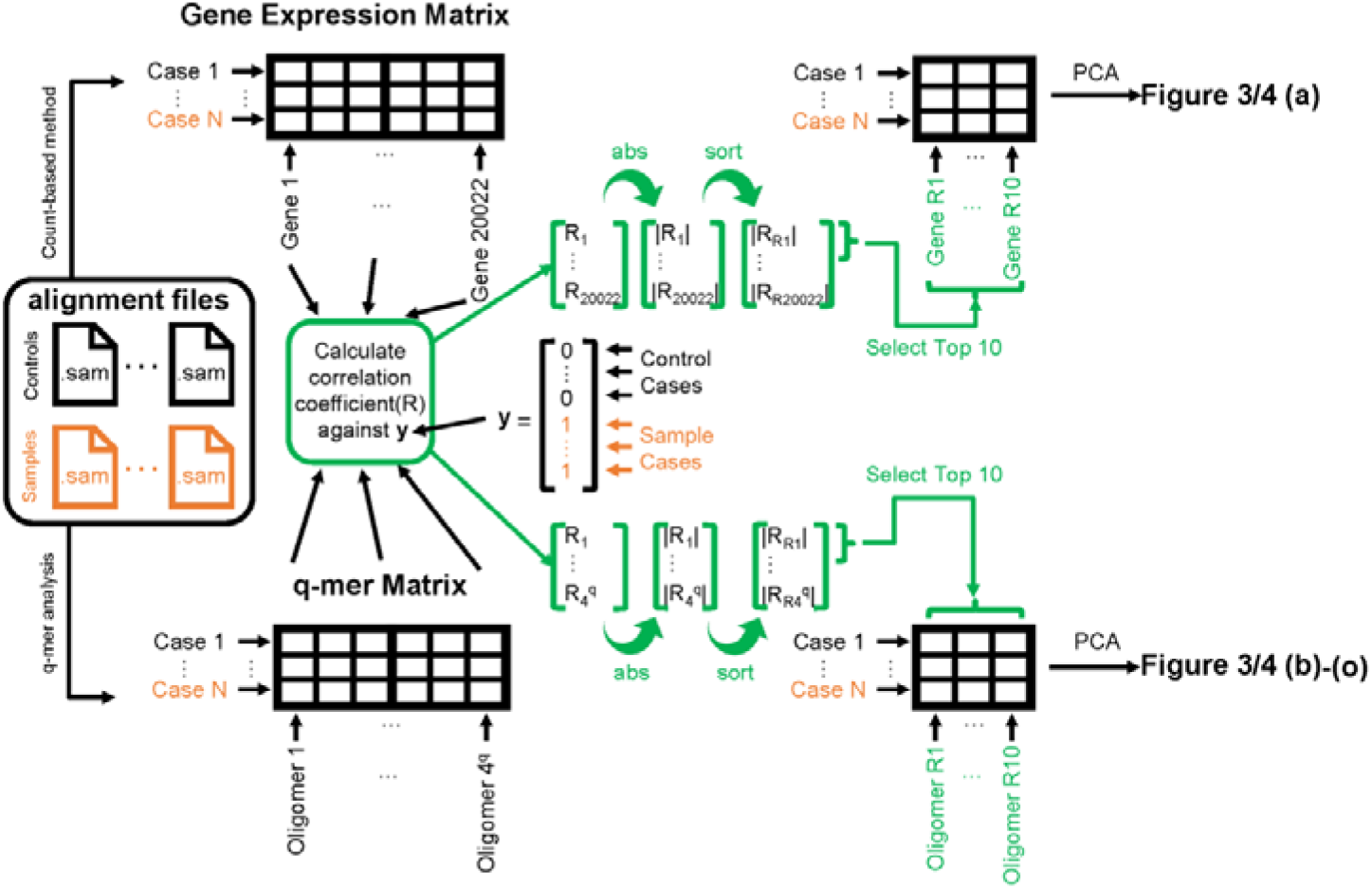
Schematic image of steps to apply principal component analysis (PCA) and draw Figures 3 and 4. The left panel shows the alignment files (black for healthy control cases and orange for cocaine-addicted or childhood-onset schizophrenia cases). The exact entity of the alignment file is a .sam file. These alignment files are converted to gene expression matrices (top-left table) by the count-based method or q-mer matrices (bottom-left table) by q-mer analysis. The correlation coefficient “R” is calculated against the ***y*** vector for each column in the table (center). The 10 columns on the top-right or bottom-right table are selected based on the size of the absolute value of “R” (indicated in green). Figures 3 and 4 were drawn based on the results of the PCA of the top-right table or bottom-left table.

Briefly, we first summarized these RNA-Seq datasets using a count-based method and obtained gene expression matrices with the sizes 36 × 20,022 or 38 × 20,022 for the first and second studies, respectively (Figure 2, top-left table). Then, after selecting 10 genes that showed the top 10 magnitudes of the correlation coefficient against the y vector (Figure 2, center), we decomposed the resulting matrix using PCA and plotted principal component (PC) 1 and PC2 for each sample (Figure 2, top-right table). The resulting plot was unable to distinguish the cocaine-addicted cases or the COS cases from the respective healthy controls (Figure 3(**a**) and Figure 4(**a**), respectively). On the other hand, when we summarized the RNA-Seq datasets using q-mer analysis with q values of 1 to 14 (Figure 2, bottom), the cocaine-addicted cases and the COS cases were linearly separated from the respective healthy control cases with q values of 12–14 and 14, respectively (Figure 3(**m–o**) and Figure 4(**o**)). Note that applying PCA to the gene expression matrix directly also failed to separate the cases from the controls (Figure S1). These results support the idea that q-mer analysis is able to describe the samples more accurately than other methods.

**Figure 3.**
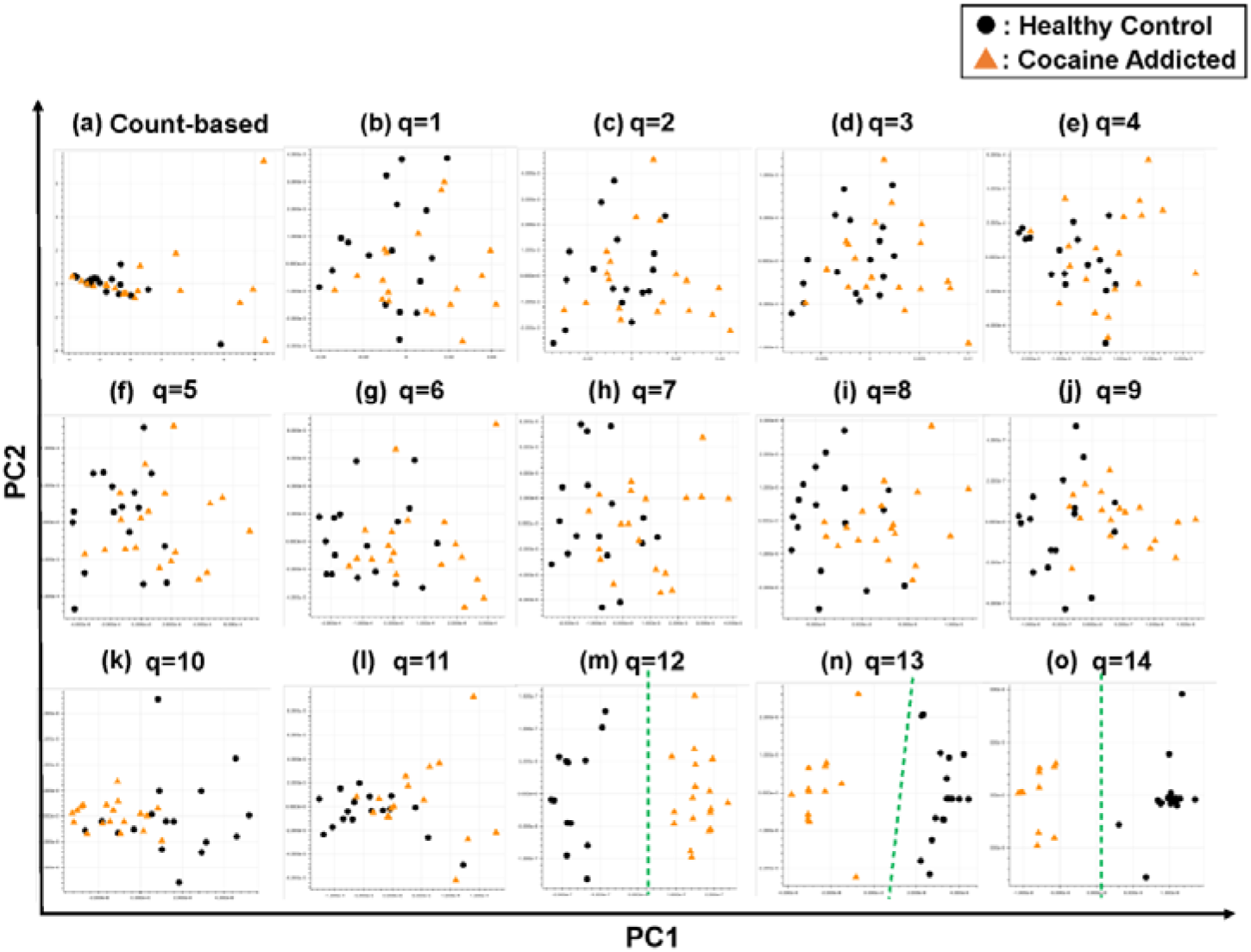
The ability of the matrices to distinguish healthy controls from the cocaine-addicted cases. The distribution of the healthy control cases (closed black circles) and cocaine-addicted samples (closed red triangles) on the PC1/PC2 plane were plotted based on the matrix from (**a**) the count-based method and (**b**–**o**) q-mer analysis. For panels (**b**)–(**o**), the q value is shown above the panel. In panels (**m**)–(**o**), a green dotted line separates the cocaine-addicted cases from the healthy controls.

**Figure 4.**
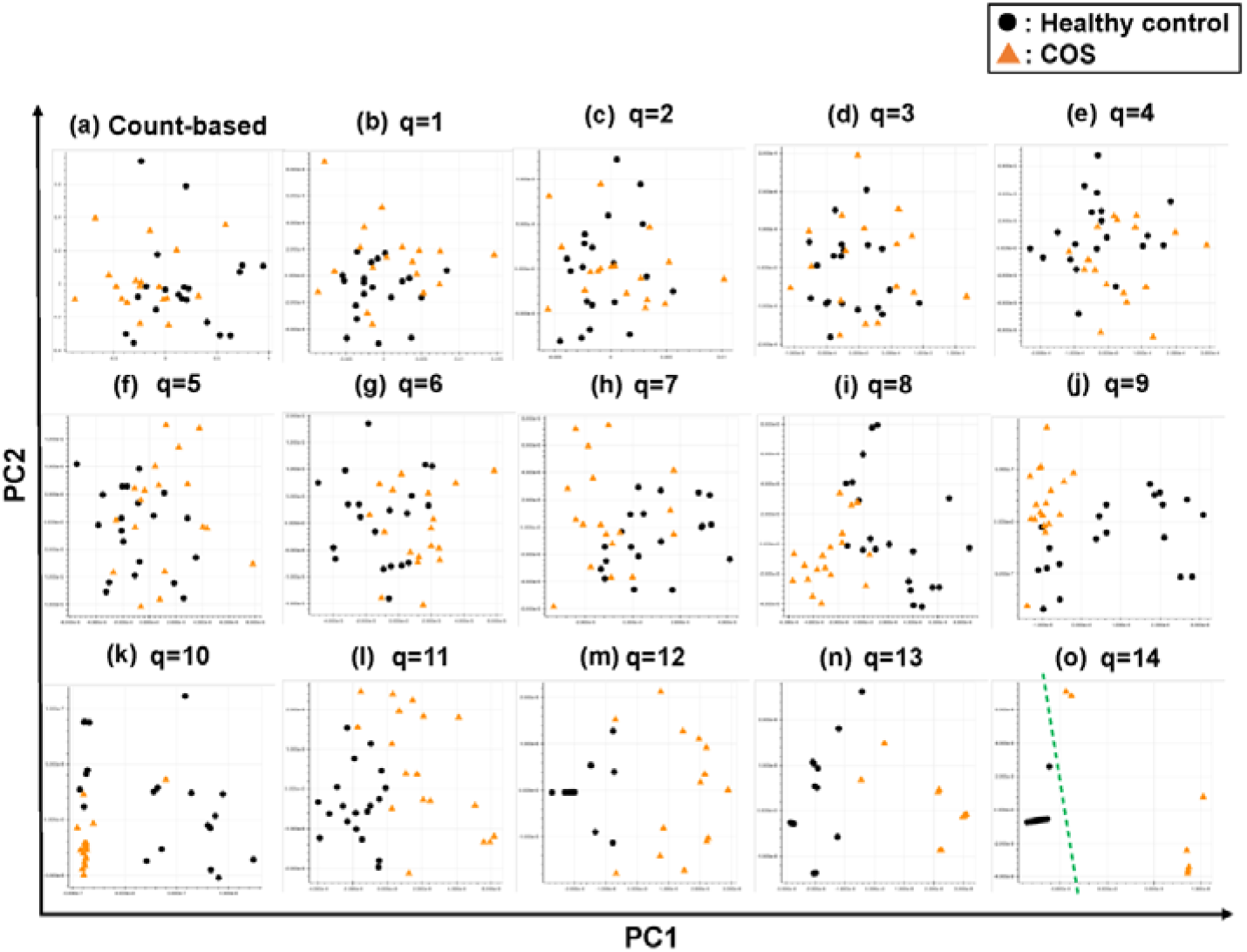
The ability of the matrices to distinguish healthy control cases from the COS cases. The distribution of the healthy control cases (closed black circles) and COS cases (closed red triangles) on the PC1/PC2 plane were plotted based on the matrix from (**a**) the count-based method and (**b)**–(**o**) the q-mer analysis. For panels (**b**)–(**o**), the q value is shown above the panel. In panel (**o**), the green dotted line separates the cocaine-addicted cases from the healthy control cases.

### 2.4. The Interpretability of q-mer Analysis

The interpretability of q-mer analysis is an important issue to confirm its utility. In other words, we next sought to investigate which biological features q-mer analysis captures to distinguish cocaine-addicted or COS cases from their respective healthy control cases.

Regarding the study of cocaine addiction, the contribution of PC1 in Figure 3(**o**) was 0.91 (Figure 5(**a**)), and all of the oligomers contributed to PC1 to the same extent (Figure 5(**b**)). These oligomers were only expressed either in the controls or in the cocaine-addicted cases and as part of one or two genes (Figure 5(**c**)). Interestingly, some of the identified genes are expressed specifically in the brain and are reported to be involved in neurological function. When examining the alignment for the region where the “ACTCGACCAAAAAT” oligomer was observed, for example, the shape of the alignment was different between the cocaine-addicted cases and the healthy controls (Figure 5(**d**)), indicating a difference in the post-transcriptional regulation between the two groups. The same feature was true for other oligomers (Figure S2). Note that the false discovery rates (FDRs) calculated based on the count-based method suggested that none of the genes in Figure 5(**c**) were significantly differentially expressed between the two groups. Furthermore, PCA of the count-based matrices only with the genes in Figure 5(**c**) failed to separate the cocaine-addicted cases from the healthy controls (Figure S3(**a**)), implying that it is essential to be able to discriminate between specific gene regions. These results suggest that q-mer analysis focuses on specific regions of each gene and is able to capture these differences in the shape of the alignments to discriminate the cocaine-addicted cases from healthy controls in this RNA-Seq dataset.

**Figure 5.**
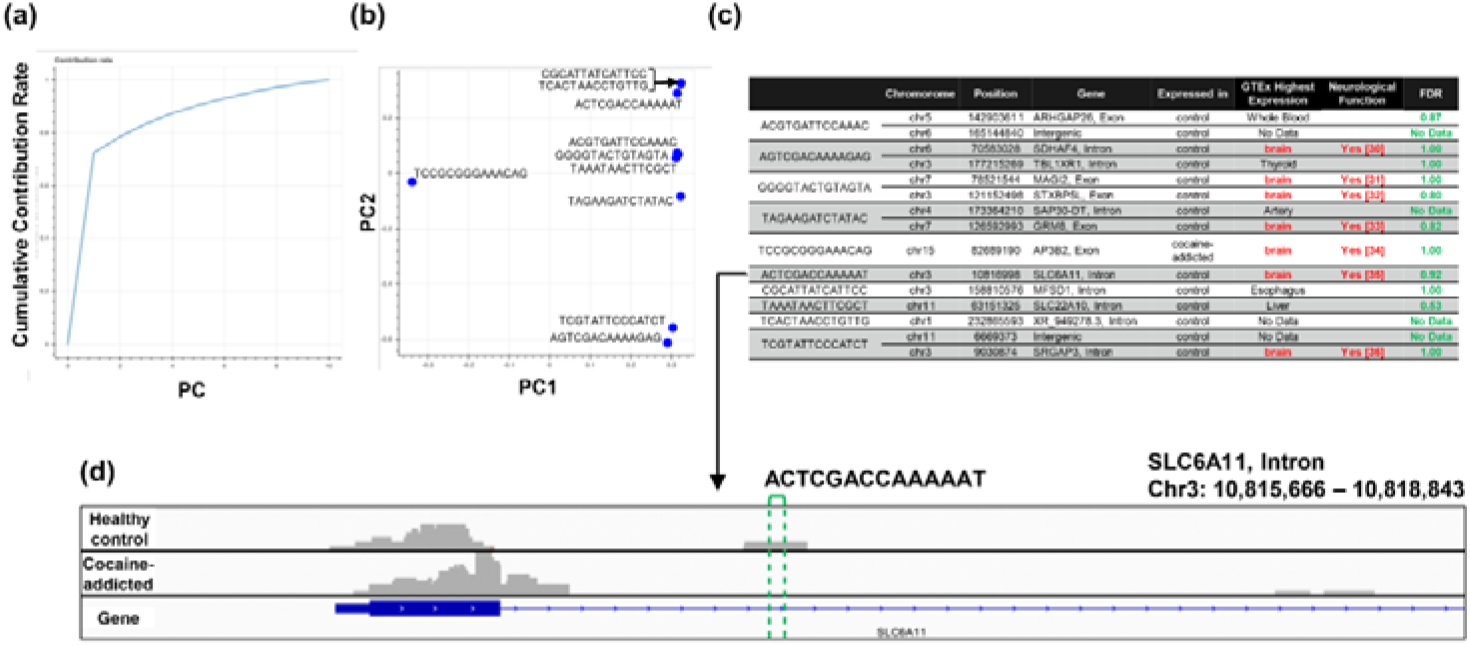
Detailed results of the q-mer analysis in the study of cocaine addiction. The cumulative contribution rate of PC1 to PC10 in the PCA of the q-mer matrix with q = 14 is shown in (**a**). The contribution rate of each oligomer for PC1 and PC2 is shown in (**b**). The position in the genome, the gene symbols for each oligomer described in the .sam files, and the false discovery rate (FDR) calculated by differential gene expression analysis based on the count-based matrix are shown in (**c**). Example alignments near the region “TAGAAGATCTATAC” are shown in (**d**). The position of the oligomer is indicated by green dotted lines.

For the study of COS, the contribution rate of PC1 in Figure 4(**o**) was 0.90 (Figure 6(**a**)), and nine oligomers contributed to PC1 to the same extent (Figure 6(**b**)). These nine oligomers were only expressed in the healthy controls (Figure 6(**c**)), six of which were observed in the same region of the TTBK2 gene (Figure 6(**c**)–(**d**)). However, there was not a clear difference in the shape of the alignment. Instead, we detected a mutation in the region. The same feature was confirmed with the other oligomers (Figure S4). These results support the idea that q-mer analysis in this RNA-Seq dataset detected mismatch information that the count-based method does not include to separate the cases from the controls.

**Figure 6.**
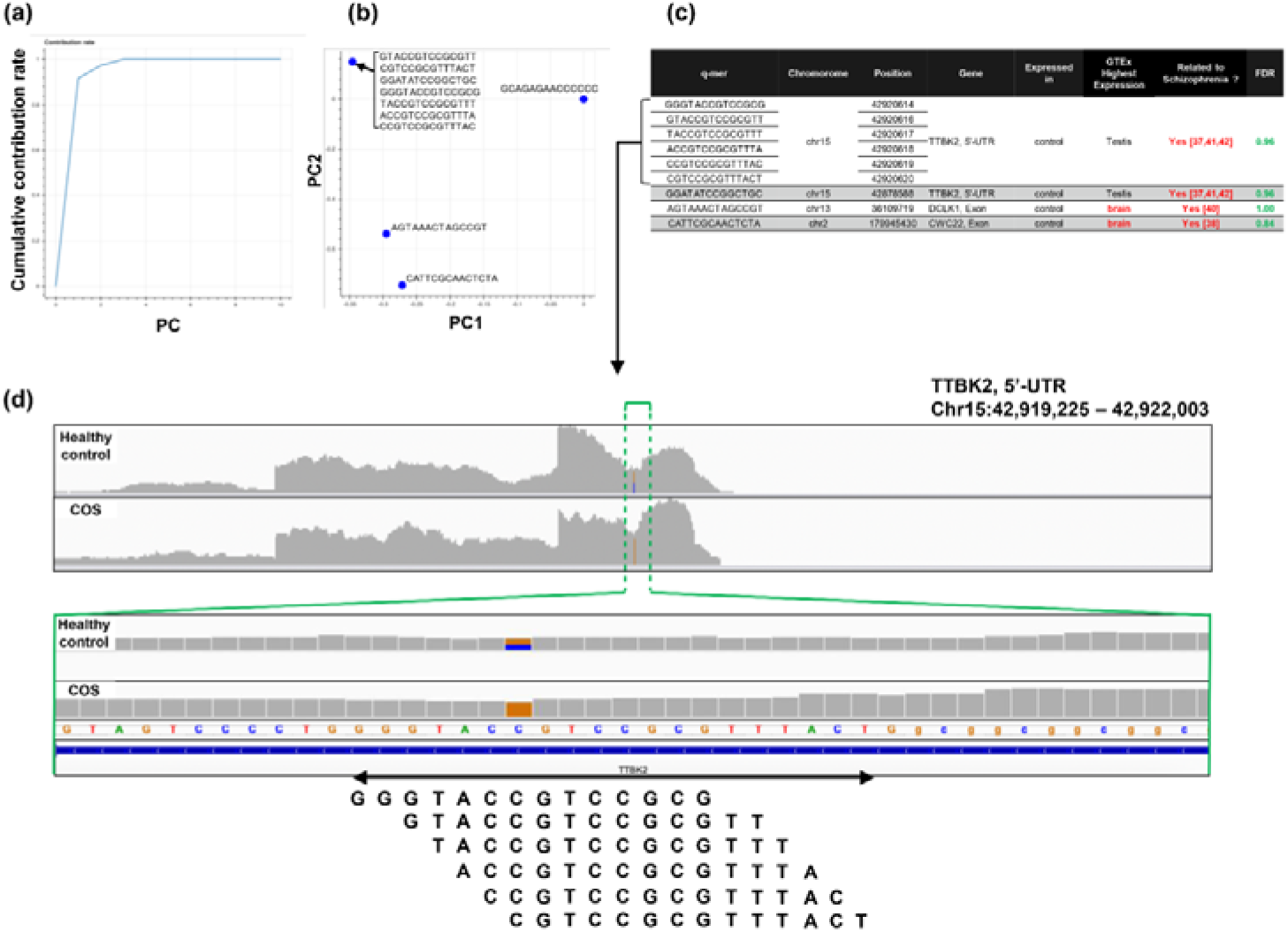
Detailed results of q-mer analysis in the COS study. The cumulative contribution rate of PC1 to PC10 in the PCA of the q-mer matrix with q = 14 is shown in (**a**). The contribution rate of each oligomer for PC1 and PC2 is shown in (**b**). The position in the genome and the gene symbols for each oligomer described in the .sam files and the FDR calculated by differential gene expression analysis based on the count-based matrix are shown in (**c**). Example alignments near the region where the six oligomers map are shown in (**d**). The position of the oligomers is indicated in green dotted lines.

## 3. Discussion

In this study, we describe a new method, q-mer analysis, that includes alignment information in RNA-Seq data analysis. q-mer analysis focuses not on the expression of whole genes but on oligomers and produces vectors with higher dimensionality than do count-based methods to summarize alignment information. This “dimensionality increment” was critical for characterizing samples so that non-supervised subgrouping was successful when analyzing the RNA-Seq datasets with large q values (Figure 3(**m**)–(**o**), Figure 4(**o**)), while the count-based method failed to distinguish between subgroups (Figure 3(**a**), Figure 4(**a**), Figure S1).

The ability of q-mer analysis to accurately characterize transcriptomics samples could be helpful when identifying the biological mechanisms underlying disorders. Currently, the diagnosis of most psychiatric or neurological diseases, such as major depressive disorder, bipolar disorder, autism spectrum disorder, and attention-deficit hyperactivity disorder, depends on behavioral and symptomatic characterization. However, q-mer analysis could help to define these disorders using q-mer vectors, potentially revealing unique characteristics that could be used for diagnosis.

q-mer analysis cannot provide a clear description of the detailed biological mechanisms that explain the differences between samples and controls; however, q-mer analysis can offer hypotheses for further investigation by capturing differences in post-transcriptional regulation or expressed mutations. Through q-mer analysis, we detected differentially expressed candidate genes in the study of cocaine addiction (Figure 5(c)). These genes are not highlighted in the original study because of the high FDR. Since q-mer analysis indicated that the post-transcriptional regulation of these genes were different between cases and controls, subsequent studies should quantify the protein levels of the genes in Figure 5(c). Interestingly, most of the genes in Figure 5(c) were specifically expressed in the brain [29] and were functionally related to neurogenesis [30–36]. In the COS study, we identified mutations as discriminators between samples from COS cases and controls. Surprisingly, all of these mutations are novel [37–40]. Although many small nucleaotide polymorphisms related to COS have been reported [37–40], the mutations we identified are worth additional follow-up studies because the genes carrying these mutation were expressed and are related to COS [37,38,40–42].

Importantly, since q-mer analysis does not require any additional experiments other than the RNA-Seq dataset, scientists can re-mine existing RNA-Seq data and may find new results. For example, studies could immediately apply q-mer analysis to publically available RNA-Seq data [43–48]. However, the aligner, such as bowtie [49], bowtie2 [50], BWA [51,52], STAR [53], and HISAT2 [54,55], that is used must be considered; use of different aligner programs from study to study may produce different results. If the alignment data are different, then q-mer analysis results may differ as well.

Furthermore, RNA-Seq data derived from libraries constructed using the poly-A method should be avoided because this method only captures the 3′-end of each mRNA transcript. Thus, this method does not capture alignment information from the 5′-end and is thus not appropriate for q-mer analysis. In addition, currently, we do not have a valid statistical method to quantify the impact of q-mer analysis. In this study, we selected the top 10 oligomers that showed the highest correlation coefficients and observed linear separation of case and control samples using PCA. However, ideally, significant oligomers should be identified statistically. Investigations of the probability distribution of q-mer results remains to be addressed in future studies.

The dimension required to express alignment information of RNA-Seq data was estimated at least 4^9^ (262,144; Table 1). Furthermore, to identify differences among the case samples and controls accurately, the required dimension may be 4^14^ (268,435,456) or larger. Thus, the dimensionality of RNA-Seq data potentially has approximately 10,000 times the number of genes than are found in *H. sapiens*. However, some reports say that the affective dimension of the transcriptome should be far less than the total number of genes it contains [56–58]. This contradiction may be because those studies only investigated ideal situations. In actual cases or conditions that have not yet been examined, such as in neurological or psychiatric disorders in *H. sapiens*, the transcriptome could have a large number of dimensions owing to the complexity of the brain and the etiology of these diseases.

The high dimensional nature of RNA-Seq data may be its advantage. Recently, scientists have attempted to describe sample conditions by combining multi-omics data to obtain additional explanatory variables. However, this approach is costly and hard to comprehend. Combining RNA-Seq experiments with q-mer analysis may be sufficient to describe samples because the dimensionality of RNA-Seq is much higher than that derived from multi-omics approaches. In the future, q-mer analysis may be the new standard rather than omics-related methods.

We suggest that there might be a limit for gene-level characterization in understanding complicated samples such as those derived from neurological and psychiatric disorders because networks and pathways often share similarities despite different disorders in each study [27]. The number of parameters (i.e., the number of genes) is probably not sufficient to explain these disorders. In contrast, q-mer analysis provides many parameters by focusing on oligomers and can separate samples if the number of samples and the q value are large enough. Further, q-mer analysis can identify candidate genes and biological mechanisms underlying differences between samples. Therefore, we propose using q-mer analysis to increase dimensionality, to identify novel mechanistic hypotheses based on differences in oligomers between different conditions, and to study underlying biology. In conclusion, differential transcriptomics based on q-mer analysis can provide novel data for clinical studies, diagnosis, prognosis, and identification of new genetic markers for diseases.

## 4. Materials and Methods

### 4.1. Criteria for Choosing Example RNA-Seq Datasets

Example RNA-Seq datasets were selected based on five criteria: (1) the dataset was related to neurological or psychiatric disorders, (2) the raw sequence data were available, (3) the RNA-Seq libraries were not constructed using the poly-A method, (4) more than 20 RNA-Seq samples were available, and (5) the RNA-Seq data were from *H. sapiens*.

### 4.2. Example RNA-Seq Datasets

Two RNA-Seq datasets were chosen based on the criteria mentioned above: GEO accession numbers GSE99349 [27] and GSE106589 [28]. The first dataset, GSE99349, contained 36 fastq files: 17 from healthy controls and 19 from cocaine-addicted cases. All 36 fastq files were used for q-mer analysis. The second dataset, GSE106589, contained 94 fastq files: 20 from hiPSCs-NPCs derived from the healthy control group, 20 from total neurons derived from the same control group, 18 from hiPSCs-NPCs derived from 14 individuals with COS, 18 from total neurons derived from the same COS group, and 18 were excluded [28]. Out of 94 files, only the 38 hiPSCs-NPC fastq files were used in order to focus our analyses on a single cell type [28].

### 4.3. Count-based Matrices and Differential Gene Expression Analysis

The 36 or 38 fastq files from the cocaine addiction study or the COS study, respectively, were preprocessed by clipping Illumina adapter sequences using Trimmomatic v.0.39 [59] and were aligned to the human genome sequence (GenBank assembly accession: GCA_000001405.28) using HISAT2 v.2.2.1 [55]. Then, the read counts for each gene were calculated by featureCounts v.2.0.1 [60] using the human gene feature file (INSDC Assembly: GCA_000001405.28, Database version: 103.38) as a reference. Finally, the resulting 36 and 38 count data files were aligned with each other to obtain the count-based matrices. The sizes of the matrices were 36 × 20,022 and 38 × 20,022, respectively. Differential gene expression was performed to obtain the FDR as shown in Figures 5(c) and 6(c), and the edgeR library [61] was applied to the count-based matrices.

### 4.4. q-mer Vectors and q-mer Matrices

First, 4^q^-dimension vectors were produced by counting the frequency of each 4^q^ kind of q-length oligomer in each .sam file. Next, the count of each oligomer was normalized by the frequency of the oligomer in the transcriptome. Then, the data were further normalized so that the sum of the elements in each vector equaled 1. Finally, the resulting 4^q^ dimension vectors were defined as the q-mer vectors. To obtain the q-mer matrices, the 36 or 38 q-mer vectors were aligned with each other. The sizes of the matrices were 36 × 4^q^ and 38 × 4^q^. Reads that were not uniquely mapped or those that contained the character “N” were skipped. The code for producing q-mer vectors from .sam files is publicly available through GitHub: https://github.com/tatsumashoji/qmer.

### 4.5. Decomposition of the Matrices

To decompose the matrices and plot the 36 or 38 cases onto a two-dimensional plane, PCA was implemented with “scikit-learn (0.24.1)” [62]. Briefly, 10 columns were selected from each matrix. These 10 columns had the top 10 highest correlation coefficients against the y vector, where the element is 0 if the corresponding case is the healthy control and 1 if otherwise. Then, the matrix with a size of 36 × 10 was decomposed using PCA. Finally, PC1 and PC2 for each case were plotted onto a two-dimensional plane. The library “scikit-learn (0.24.1)” was used for the detailed analysis of the PCA plots shown in Figure 5(**a, b**) and Figure 6(**a, b**). To reproduce our results, we have provided all code in a Jupyter notebook at https://github.com/tatsumashoji/qmer.

### 4.6. Identification of the Genome Position of the Oligomers

Not all of the cases in the cocaine-addicted group, the COS group, or the healthy control group expressed each oligomer. The genome position for each oligomer as shown in Figure 5(c) and Figure 6(c) was defined if the oligomer was expressed at the same genome position among more than four cases.

## 5. Conclusions

We introduced q-mer analysis, a generalized method for analyzing RNA-Seq data that includes alignment information and demonstrated that this alignment information was essential to characterize the samples appropriately. Aspects that q-mer analysis can more correctly represent than count-based approaches could be helpful for studies regarding gene expression. In the future, combining RNA-Seq with q-mer analysis could help identify biologically relevant features in transcriptomics data.

## Supplementary Materials

The following are available online at www.mdpi.com/xxx/s1, Figure S1: PCA of the gene expression table without gene selection; Figure S2: Example alignments near the region where the oligomers in Figure 5(**c**) are mapped; Figure S4: Example alignments near the region where the oligomers in Figure 6(**c**) are mapped; Figure S3: PCA of the gene expression table with the genes in Figure 5(**c**) and in Figure 6(**c**).

## Author Contributions

Conceptualization, S.T.; methodology, S.T.; software, S.T.; validation, S.Y.; formal analysis, S.T.; investigation, S.T.; resources, S.T. and S.Y.; data curation, S.T.; writing—original draft preparation, S.T.; writing—review and editing, S.T. and S.Y.; visualization, S.T.; supervision, S.Y.; project administration, S.T.; funding acquisition, S.Y. All authors have read and agreed to the published version of the manuscript.

## Institutional Review Board Statement

Institutional Review Board approval are not applicable as this study does not include human or animals.

## Informed Consent Statement

Informed Consent is not applicable as this study does not include human.

## Data Availability Statement

Publicly available datasets were analyzed in this study. This data can be found here: https://www.ncbi.nlm.nih.gov/geo/query/acc.cgi?acc=GSE99349/GSE99349 and https://www.ncbi.nlm.nih.gov/geo/query/acc.cgi?acc=GSE106589/GSE106589.

## Acknowledgments

Not applicable.

