## Supplementary file for "q-mer analysis: a generalized method for analyzing RNA-Seq data"

**Supplementary Figures**

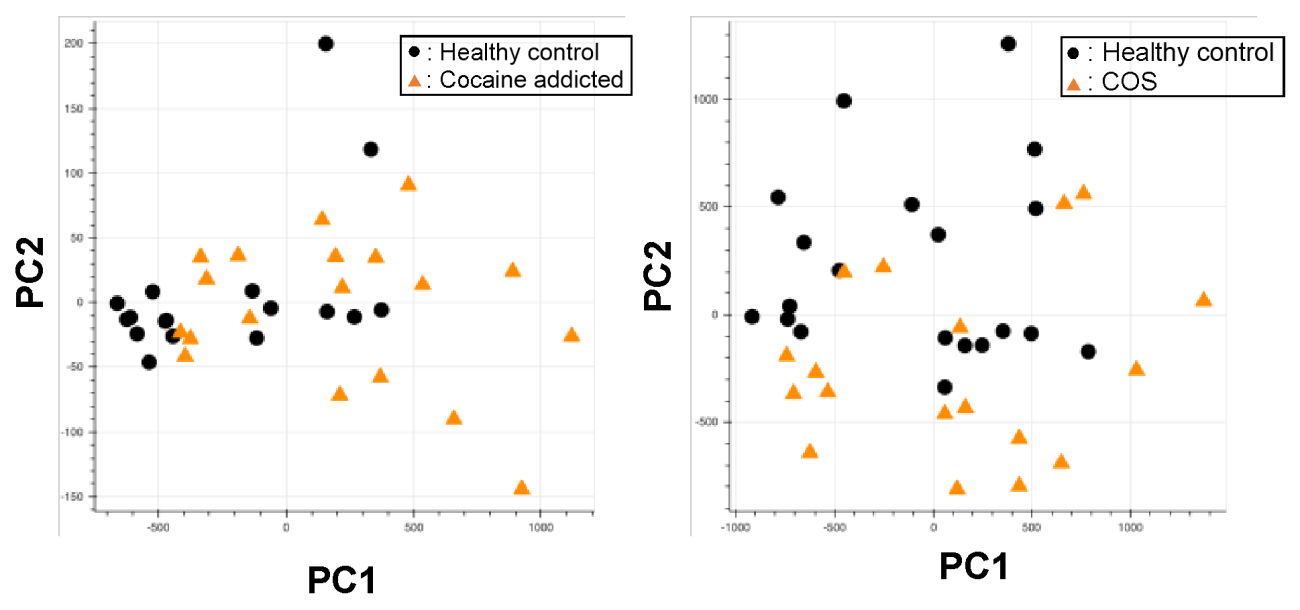

**Figure S1.** PCA of the gene expression table without gene selection for the study of cocaine addiction (**a**) and childhood-onset schizophrenia (COS) (**b**).

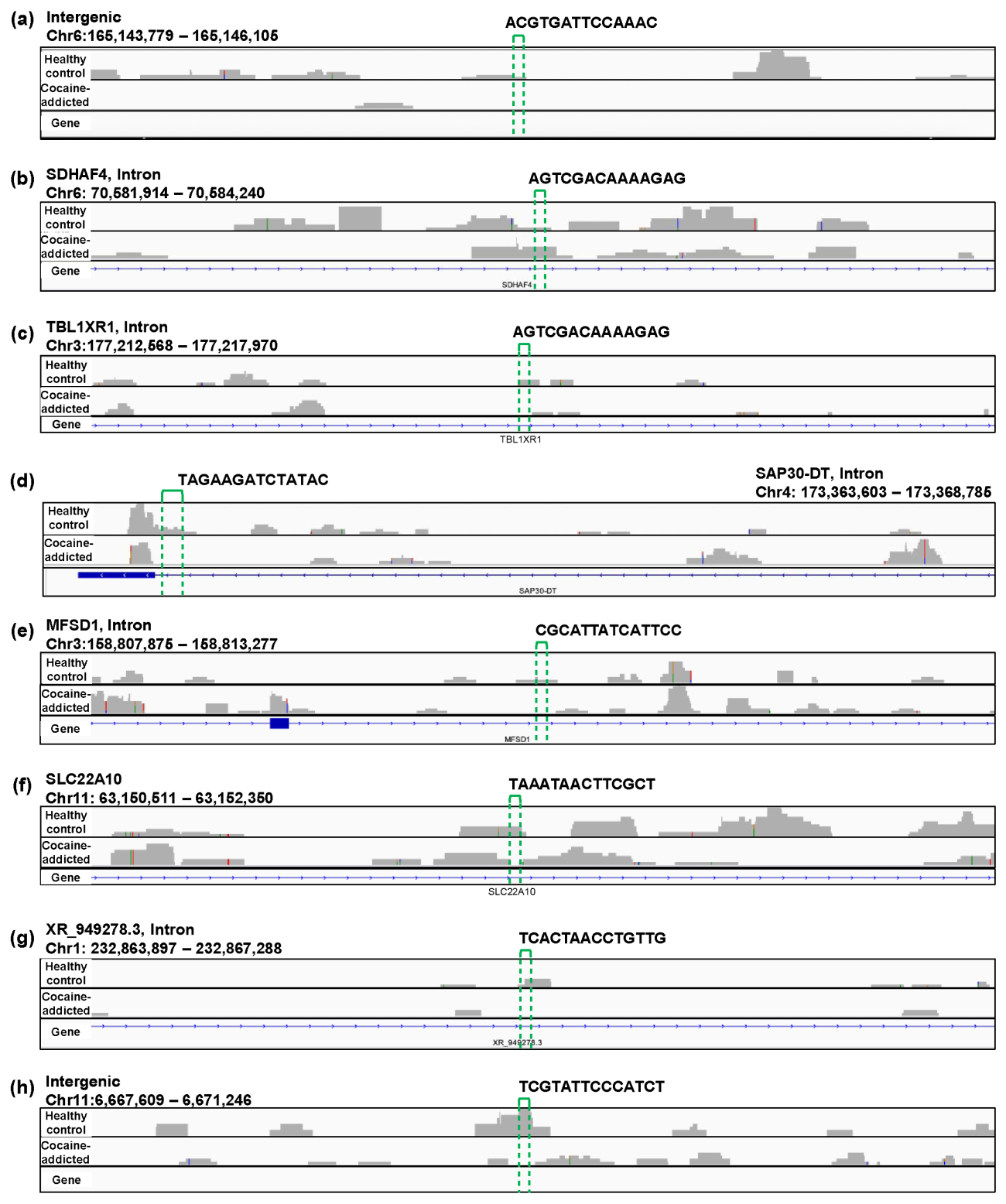

**Figure S2.** Example alignments near the region where the oligomers in Figure 5(**c**) are mapped. The positions of the oligomers are indicated by green dotted lines.

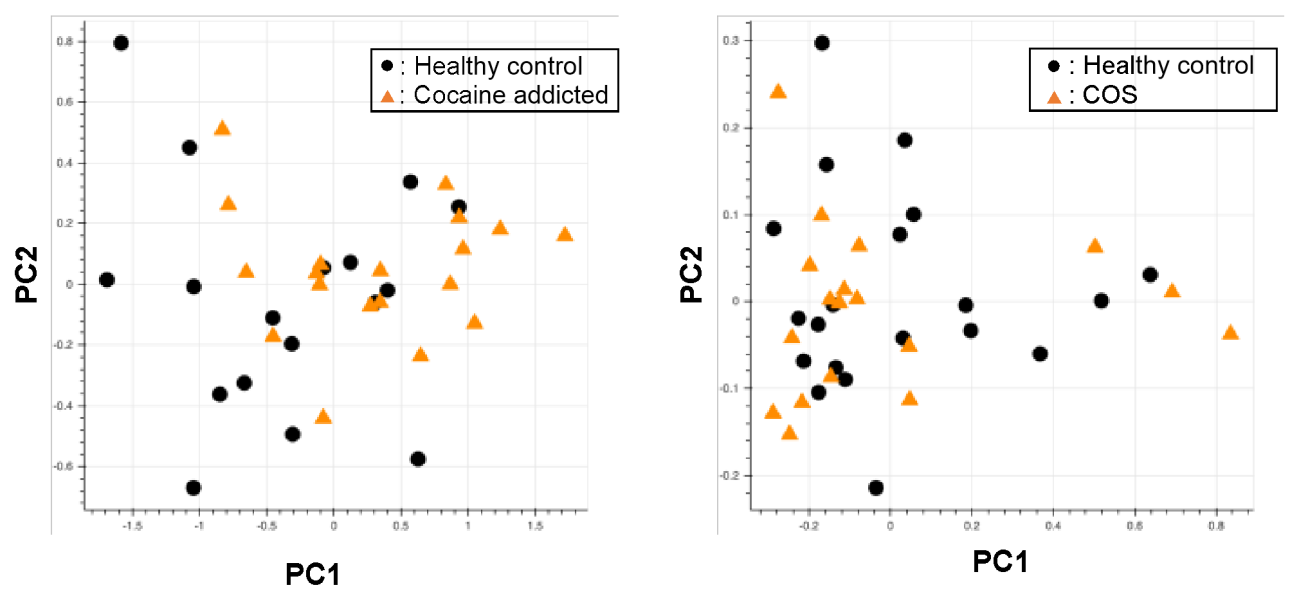

**Figure S3.** PCA of the gene expression table with the genes in Figure 5(**c**) and in Figure 6(**c**).

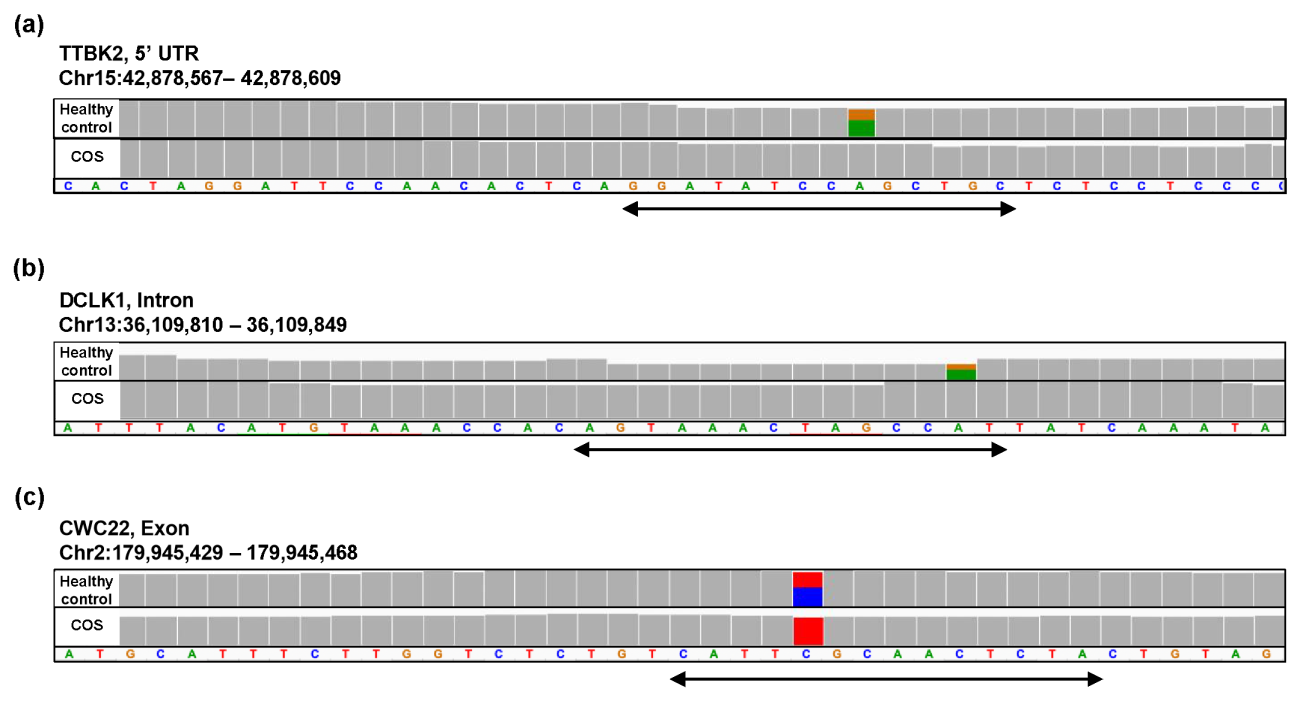
 **Figure S4.** Example alignments near the region where the oligomers in Figure 6(**c**) are mapped. The position of the oligomers is indicated by the arrow.
